# Profiling of *N*^6^-methyladenosine dynamics indicates regulation of oyster development by m^6^A-RNA epitranscriptomes

**DOI:** 10.1101/2021.08.30.458180

**Authors:** Lorane Le Franc, Bruno Petton, Pascal Favrel, Guillaume Rivière

## Abstract

The *N*^6^-methylation of RNA adenosines (m^6^A) is an important regulator of gene expression with critical implications in vertebrate and insect development. However, the developmental significance of epitranscriptomes in lophotrochozoan organisms remains unknown.

Using MeRIP-seq, we generated transcriptome-wide m^6^A-RNA methylomes covering the whole development of the oyster from oocytes to juveniles. Oyster RNA classes display specific m^6^A signatures, with mRNAs and lncRNAs exhibiting distinct profiles and being highly methylated compared to transposon transcripts. Epitranscriptomes are dynamic and correspond to chronological steps of development (cleavage, gastrulation, organogenesis and metamorphosis), with a minimal mRNA and lncRNA methylation at the morula stage followed by a global increase. mRNA m^6^A levels are correlated to transcript levels and shifts in methyladenine profiles correspond to expression kinetics. Differentially methylated transcripts cluster according to embryo-larval stages and bear the corresponding developmental functions (cell division, signal transduction, morphogenesis and cell differentiation). The m^6^A level of transposon transcripts is also regulated and peaks during the gastrulation.

We demonstrate that m^6^A-RNA methylomes are dynamic and associated to gene expression regulation during oyster development. The putative epitranscriptome implication in the cleavage, maternal-to-zygotic transition and cell differentiation in a lophotrochozoan model brings new insights into the control and evolution of developmental processes

## Introduction

The success of development of metazoan organisms is conditioned by the precise temporo-spatial regulation of gene expression. The *N*^6^-methylation of RNA adenosines (*N*^6^-methyladenosine, m^6^A) has recently emerged as a critical layer of the gene expression regulatory network. Indeed, in vertebrates m^6^A-RNA methylation is required for the proper regulation of developmental processes such as cell differentiation (Zhang et al. 2020a; Ping et al. 2014; Zhang et al. 2017), X chromosome inactivation (Coker et al. 2020; Patil et al. 2016), maternal-to-zygotic transition (MZT) (Zhao et al. 2017; Sui et al. 2020) and neurogenesis (Yoon et al. 2017; Wu et al. 2018; Edens et al. 2019). Besides, the invalidation of enzymes which deposit m^6^A on RNA is lethal during the early development (Mansour et al. 2015; Wang et al. 2014b).

The m^6^A-RNA methylation is regulated by an enzymatic machinery comprising writers and erasers. The METTL3/ METTL14/WTAP writer core complex (Yang et al. 2018) deposits methyl marks at the consensus sequence DRACH (D=A/G/T, R=A/G, H=A/C/T) (Ke et al. 2015; Meyer et al. 2012; Batista et al. 2014; Xiao et al. 2019). Erasers such as ALKBH5 (Zheng et al. 2013) and the controversial FTO (Jia et al. 2011) remove the methylation. The combined action of m^6^A writers and erasers makes m^6^A profiles highly dynamic during the development of investigated species (Wang et al. 2021; Zhang et al. 2018; Sui et al. 2020).The biological effects of RNA methylation are mediated by reader proteins able to bind m^6^A. For example, the YTH protein family of readers is involved in RNA stability, translation level (Wang et al. 2015, 2014a; Hsu et al. 2017; Shi et al. 2017), splicing (Xiao et al. 2016) and nuclear export (Roundtree et al. 2017). Other readers like hnRNPA2B1 (Alarcón et al. 2015) and IGF2BP (Huang et al. 2018) and Prrc2a (Wu et al. 2018) mediate m^6^A influence on RNA stability, and eIF3a guide cap-independent translation of m^6^A-RNAs (Meyer et al. 2015). This whole machinery allows guiding the methylation on specific RNA targets and modulates their cellular fate. As a result, m^6^A is a chemical modification that influences gene expression without modifying the associated transcript nucleic acid sequence and is therefore referred to as ‘epitranscriptomic’ (Saletore et al. 2012).

Many RNA classes are subjected to m^6^A methylation, which is the most abundant internal modification in eukaryotic messenger RNAs (mRNAs). *N*^6^-methyladenosines in mRNAs are mostly found in the 3’ untranslated transcribed region (UTR) at the vicinity of the stop codon (Meyer et al. 2012; Dominissini et al. 2012; Ke et al. 2015; Linder et al. 2015). However, m^6^A is also present at lower levels in 5’-UTRs (Coots et al. 2017; Meyer et al. 2015), long internal exons (Dominissini et al. 2012; Batista et al. 2014) and introns (Louloupi et al. 2018; Xiao et al. 2019).

While the *N*^6^-methyladenosine modification is well described in mRNA targets, it also affects most non-coding RNA classes (ncRNAs) including microRNAs (miRNAs) (Alarcón et al. 2015), circular RNAs (circRNAs) (Yang et al. 2017). small nuclear (snRNAs) (Warda et al. 2017) and nucleolar RNAs (snoRNAs) (Huang et al. 2017). In vertebrates, more than 300 lncRNAs are as m^6^A-methylated (Meyer et al. 2012), a modification that alters the RNA structure towards an increased protein accessibility (Liu et al. 2017), as well as subcellular localization (Wu et al. 2019). The m^6^A-modification of lncRNAs has critical developmental outcomes, as indicated by the requirement of m^6^A-methylation of the XIST lncRNA for the transcriptional silencing of the inactivated X chromosome (Patil et al. 2016). Besides, the m^6^A in RNA encoded by transposable element genes (repeat RNAs) increases their stability and promotes chromatin compaction (Liu et al. 2020).

Aside Vertebrates, the *N*^6^-A methylation of RNA was described in a wide diversity of organisms, such as insects (Lence et al. 2016; Jiang et al. 2019; Kan et al. 2017), yeasts (Bodi et al. 2010) and plants (Luo et al. 2014). However, despite the developmental significance of m^6^A is conserved across evolution, several differences exist between animal groups. Indeed, in the fruit fly, m^6^A is mostly present in the 5’UTR of mRNAs throughout head and embryo development (Kan et al. 2017; Worpenberg et al. 2021), whereas methylation enrichment at 5’UTRs promotes cap-independent translation for transcript selection during the stress response in Vertebrates (Zhou et al. 2015; Meyer et al. 2015; Coots et al. 2017). In the silkworm *Bombyx mori*, mRNA is mostly m^6^A-methylated in coding sequences (CDS) but not UTRs, and a high methylation is associated with a high gene expression (Li et al. 2019). Besides, a higher m^6^A content and gene expression regulation is found upstream the diapause (Jiang et al. 2019), a development phase reflective of a highly plastic phenotype. Finally, in *Caenorhabditis elegans*, where only few actors of the m^6^A machinery are present, methylation is mostly found in rRNAs (Sendinc et al. 2020). This situation opens crucial questions about the evolution of m^6^A-related molecular pathways, target genes and developmental significance that require investigations in divergent models. However, despite being of upmost importance for our understanding of the evolution of the molecular control of developmental processes, there is a critical lack of knowledge in lophotrochozoan organisms where epitranscriptomes were not investigated to date to our knowledge.

The Pacific oyster *Crassostrea gigas* (i.e. *Magallana gigas*) is a bivalve mollusk whose great ecological and economical significance allowed its emergence as a model species within lophotrochozoan organisms. As such, an important amount of genetic, transcriptomic and epigenetic data has been generated in this model (Zhang et al. 2012; Penaloza et al. 2020; Riviere et al. 2013; Dheilly et al. 2011). Besides, the embryo-larval development of *C. gigas* is under the strong epigenetic influence of DNA methylation (Riviere et al. 2017; Sussarellu et al. 2018; Saint-Carlier and Riviere 2015; Riviere et al. 2013) and histone marks (Fellous et al. 2014, 2019). Moreover, it has recently been demonstrated that m^6^A and its complete associated machinery are conserved in the oyster, with features strongly indicating epitranscriptomic regulation of development (Le Franc et al. 2020). Furthermore, oysters develop as pelagic larvae in sea water before they metamorphose into fixed adult specimen. Therefore oyster early life stages are directly subjected to the variations of external environmental conditions such as temperature changes, UV exposure or endocrine disruptor contamination, which are described to influence m^6^A-RNA methylation in investigated species (Lu et al. 2019; Xiang et al. 2017; Cayir et al. 2019; Meyer et al. 2015; Zhou et al. 2015). These elements are highly suggestive of a functional significance of m^6^A-RNA methylation in oyster development, which remains unknown to date.

To investigate these questions, we characterized the transcriptome-wide dynamics of m^6^A-methylomes across the entire development of the pacific oyster from the oocytes to the completion of the organogenesis. We used MeRIP-seq to investigate and monitor the dynamics of m^6^A levels, localization, target RNA subsets and functional implications in a lophotrochozoan model to provide a better understanding of the evolution of developmental mechanisms and their epigenetic regulation.

## Results

### Oyster RNA classes display specific developmental m^6^A signatures

In developing oysters, m^6^A methylation affects a vast majority of RNA classes, and is detected in 91.0% of mRNAs, 65,7% of lncRNAs, 87.0% of RNA transposable elements (TEs) (retrotransposons) and 99.4% of DNA TEs. In contrast, oyster tRNA are mostly unmethylated (rRNA were depleted during library preparation and not investigated here). In addition, 35.8% of mRNAs and 11.1% of lncRNAs are significantly m^6^A-enriched over background levels (**Figure 1A**). The methylation pattern depends on the RNA class considered (ks test *p*<7,55.10^−11^). While m^6^A in mRNA is found from the 5’- to the 3’UTR, and is more abundant around the stop codon, it is more randomly spread along lncRNAs (**Figure 1B**). Methylation affects adenosines located within the ‘AGUGA*C’ sequence (where * marks the modified adenosine) (**Figure 1C**). The mean methylation level differs between RNA classes and DNA TE transcripts are less methylated than mRNAs, lncRNAs and retrotransposons RNAs (**Figure 1D**). The MeRIP–seq procedure was validated by investigation of the status of five transcripts exhibiting contrasted m^6^A vs. mRNA profiles in other species (*c-myc*, *klf*, *mettl3*, *hnrnpa2b1* and *oct4*) by targeted MeRIP-qPCR. All the examined candidates displayed similar patterns between the two techniques and consistent with the literature (**Supplemental Figure S1**).

**Figure 1:**
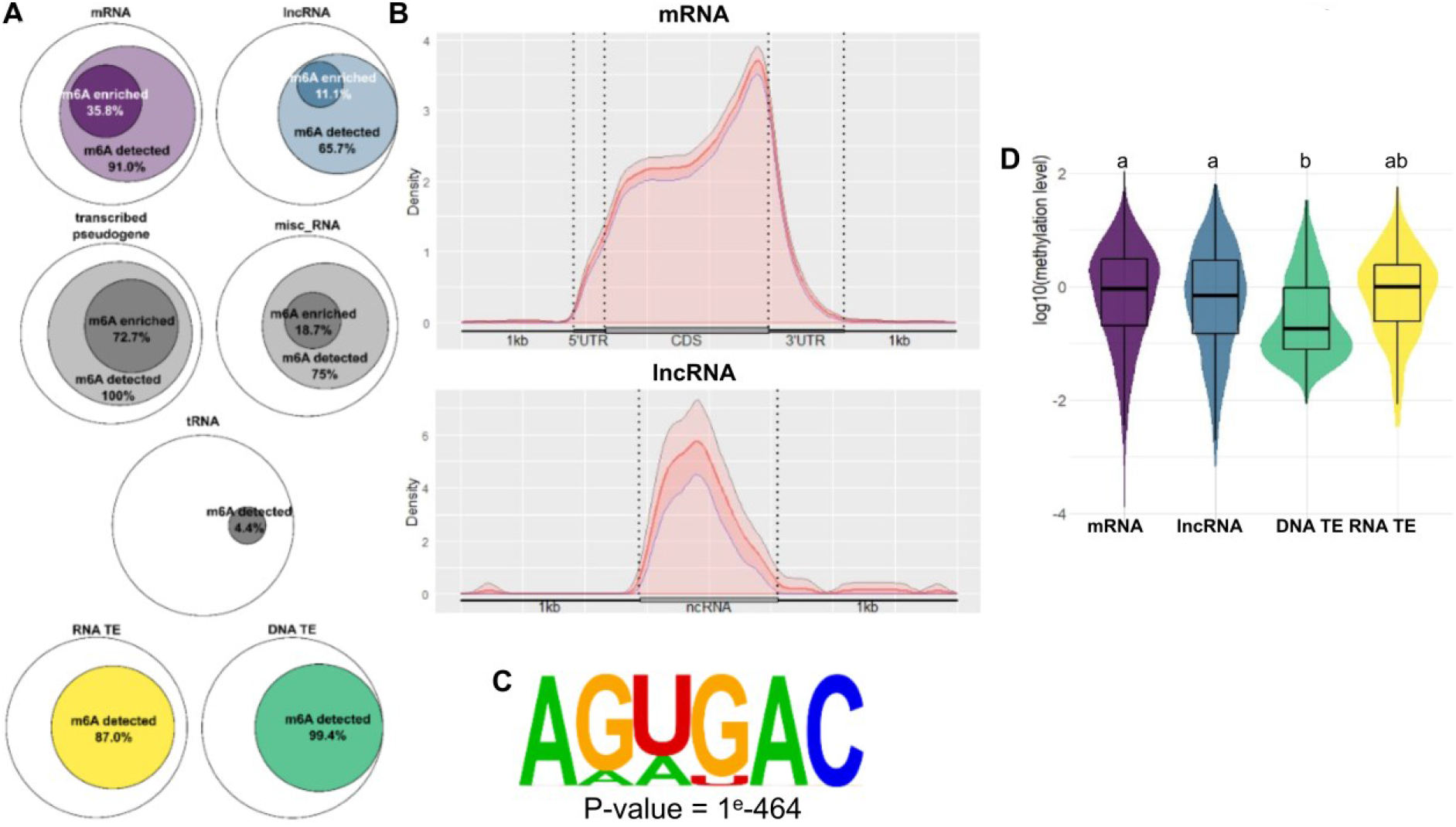
*N*^6^-methyladenosine signatures of oyster RNA classes. **A**. m^6^A distribution among RNA classes (mRNA, messenger RNA; lncRNA, long non-coding RNA; DNA TE, DNA transposable elements; RNA TE, retrotransposons; misc. RNA, miscellaneous RNA; tRNA, transfer RNA). Diagram diameter was normalized between RNA classes. The internal circles in diagrams represent the proportion of RNA displaying significant m6A methylation over background (light color) and significant m6A enrichment (dark color), respectively. **B**. m^6^A localization along mRNAs and ncRNAs (5’ to 3’). The mean m^6^A density and confidence interval are represented. UTR: untranslated transcribed region, CDS: coding sequence, kb: kilobase. **C**. Consensus sequence of the m^6^A motif in the oyster identified by HOMER. **D**. Mean methylation level of the methylated transcripts of mRNA, lncRNA, DNA TE and retrotransposons (RNA TE) classes during oyster development. Letters discriminate significantly different methylation levels (ANOVA followed by Bonferroni’s post hoc test, p<0,05).

### Oyster m^6^A epitranscriptomes are dynamic and correspond to defined steps of the development chronology

The Principal Component Analysis (PCA) of MeRIP-seq data indicates that the samples group together according to the embryo-larval stages and the samples are overall ordered chronologically along the X-axis (PC1) (**Figure 2A**). The PCA plot can be divided into four areas defining four developmental phases: cleavage (from oocytes to 2-8 cell-stage), gastrulation (blastula and gastrula stages), tissue differentiation (trochophore and D-larvae stages) and metamorphosis (pediveliger and spat stages). The morula stage is clearly individualized along the PC2, suggesting peculiar m^6^A-methylation and development features. This segregation is confirmed by the pairwise correlation matrix, which also indicates a surprising proximity of cleavage and metamorphosis epitranscriptomes (**Figure 2B**). The chronological segregation of developmental stages according to m^6^A methylation indicates that m^6^A-RNA epitranscriptomes are an important component of the developmental process.

**Figure 2:**
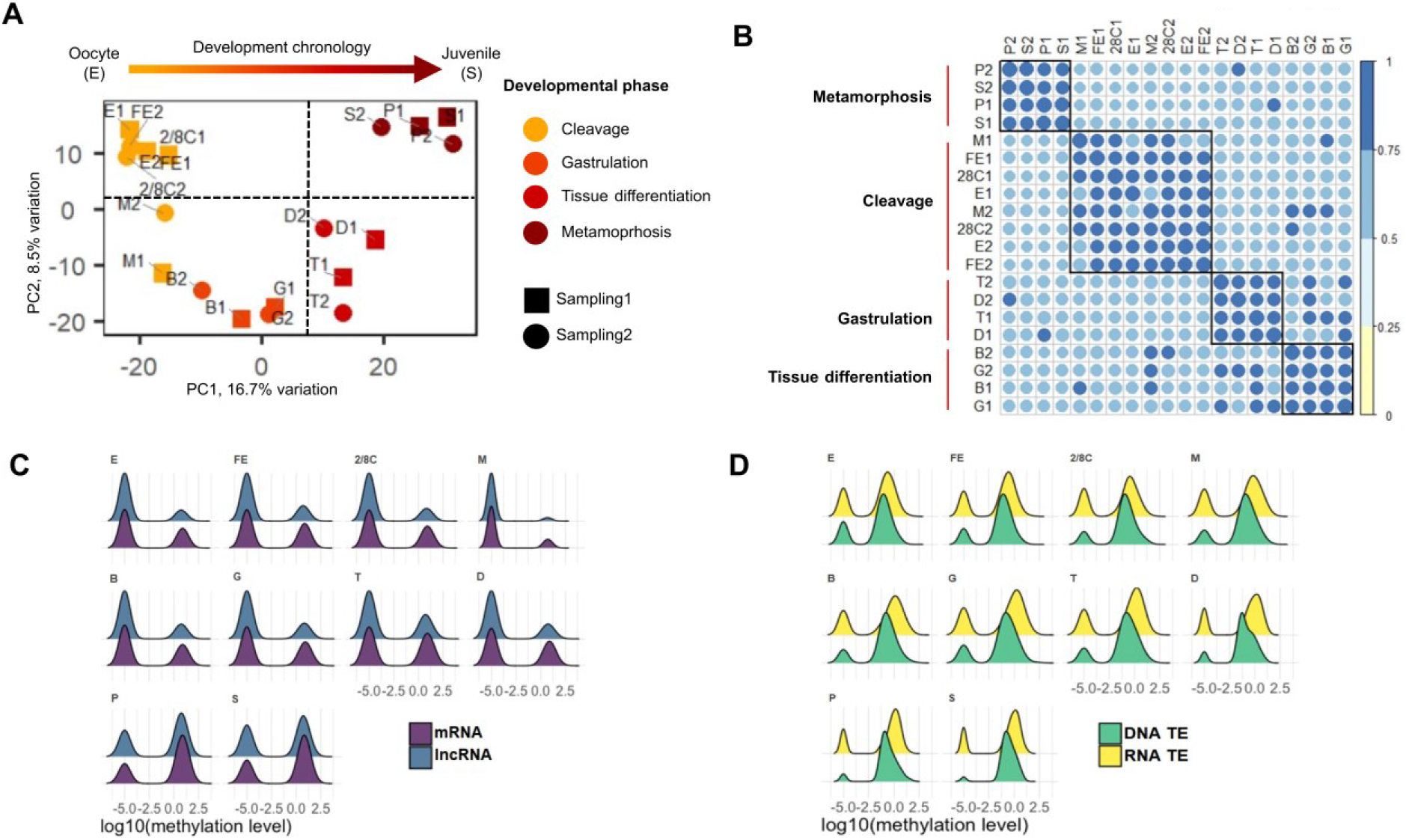
Epitranscriptome dynamics during oyster development. **A.** Principal component analysis of MeRIP-seq results. PCA plot of the first two principal components of the MeRIP-seq results for each developmental sample of the two distinct experiments. **B**. Similarity (pairwise Pearson’s correlation matrix) of m^6^A-methylation between samples based on IP/Input signal (see methods). **C**. **and D.** Dynamics of the m^6^A-methylation level of the mRNA (purple) and lncRNA (blue) (**C**); and of the DNA TE (green) and RNA TE (yellow) (**D**) transcripts. All the transcripts found m^6^A-enriched over background using MetPeak in at least one development stage are represented. Areas under the curve are normalized for each RNA class, and possible undetectable methylation at certain stages was arbitrarily affected a -5 value for representation purpose. E: Egg (oocyte), FE: Fertilized egg, 2/8C: Two to eight cell-stage, M: Morula, B: Blastula, G: Gastrula, T: Trochophore, D: D-larva, P: Pediveliger, S: Spat. mRNA (messenger RNA), lncRNA (long non-coding RNA), DNA TE (DNA transposable element), RNA TE (RNA transposable element).

The m^6^A epitranscriptomes are dynamic during oyster development. There is an important drop in the number of methylated transcripts at the morula stage followed by a general shift towards increased m^6^A-methylated mRNAs and lncRNAs onwards, which is especially marked at the late development stages (i.e. pediveliger and spat stages) (**Figure 2C**). The same tendency is also observed, although less importantly, for TE RNAs, which unmethylated population becomes reduced in the late developmental phases (**Figure 2D**).

### mRNA m^6^A profiles are regulated and associated to expression level and kinetics throughout oyster development

The mRNA m^6^A methylation is highly dynamic during oyster development. Indeed, among the significantly methylated mRNAs (ANOVA *p*<0.05, n= 8404), 17.8% of m^6^A-mRNA (n=1494) display a significant variation in their m^6^A level (**Figure 3A**), and this m^6^A level is significantly correlated to their expression level (**Figure 3B**). The differentially methylated (DM) m^6^A-mRNAs define 4 clusters according to their methylation (and expression) profiles, corresponding to the developmental steps defined previously. The cluster 1 includes mRNAs with a strong methylation and transcript content in the early stages (up to the morula stage) which decreases afterwards. By contrast, the mRNAs within the three other clusters display poor methylation during early stages, which strongly increases from the gastrulation (cluster 2), the trochophore stage (cluster 3) and the pediveliger stage (cluster4) and remain strongly methylated afterwards. Overall, there is a marked decrease of m^6^A methylation at the morula stage.

**Figure 3:**
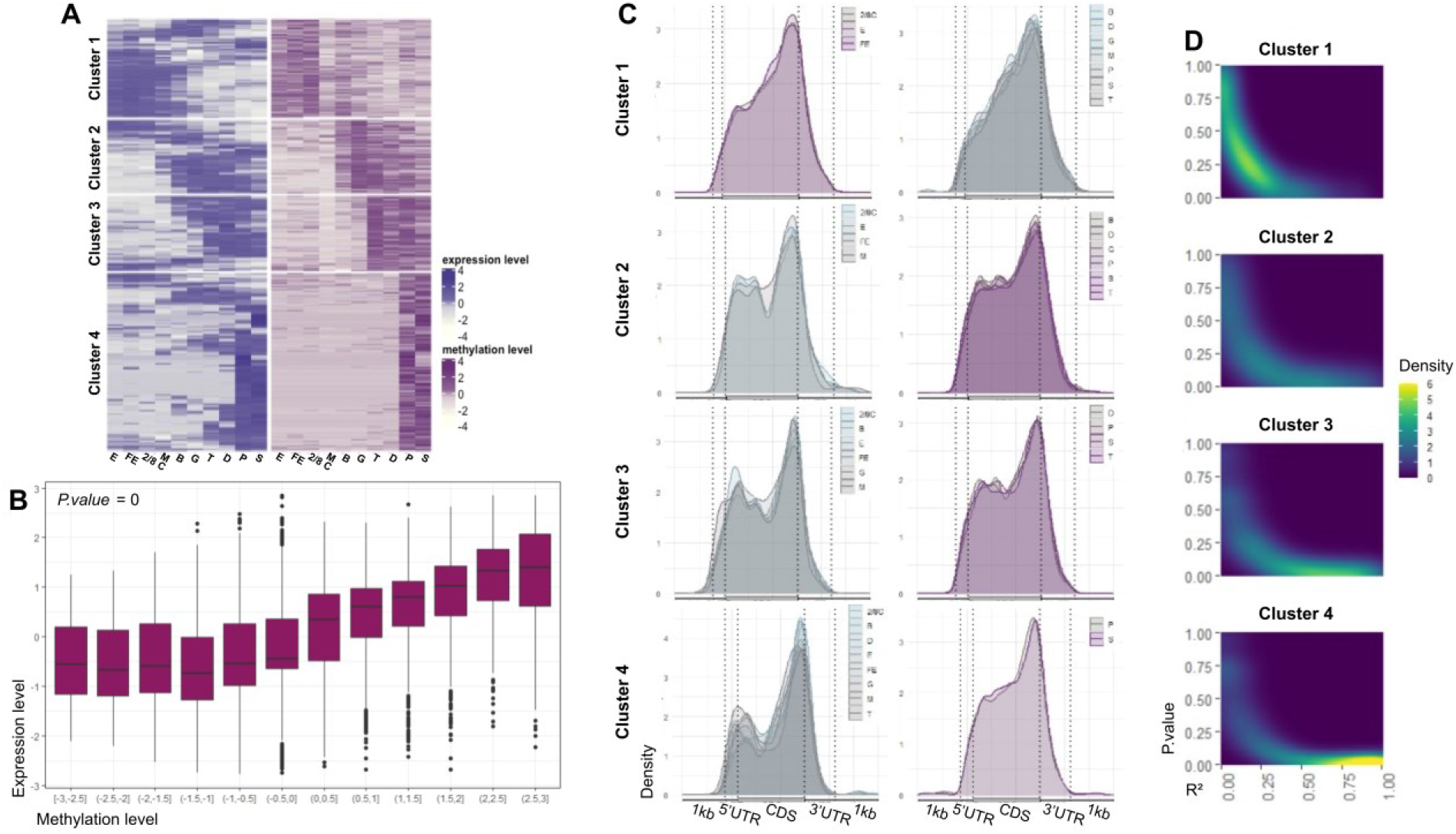
Dynamics of mRNA expression and m^6^A methylation during oyster development. **A.** Normalized expression (TPM, purple) and m^6^A content (IP/Input, pink) of significantly differentially methylated mRNA genes during oyster development (n=1494). The development stages are indicated below. **B**. Correlation between expression (Y axis, TPM) and methylation (X axis, IP/Input, divided into twelve quantiles). **C**. Methylation profiles of mRNAs displaying high (pink) or low (grey) expression within each cluster. The development stages in high and low expression groups are indicated (mean m^6^A density; confidence intervals were omitted for clarity). **D**. Correlation of expression vs. methylation dynamics throughout development. The linear correlation of the methylation variation vs. expression variation was assessed for each gene across oyster development per cluster and the results are given as surface plots (R-square: X-axis, *p*-value: Y axis, density of genes: Z axis, blue to yellow). Yellow color in the right bottom corner (i.e. *p* value<0;05 and *R^2^*>0.5) indicates strong correlation. E: Egg (oocyte), FE: Fertilized egg, 2/8C: Two to eight cell-stage, M: Morula, B: Blastula, G: Gastrula, T: Trochophore, D: D-larva, P: Pediveliger, S: Spat.

Besides the level of methylation, the localization of m^6^A within mRNAs can also vary between clusters (ks test, 0.31<*p*<5.11.10^−12^ depending on cluster pairwise comparison). Indeed, cluster 1 mRNAs, which are essentially maternal mRNAs, are methylated mostly around their stop codon in oocytes and thereafter. By contrast, mRNAs within the three other clusters, that are essentially expressed from the zygotic genome, display a marked biphasic profile, with an increased 5’ m^6^A content at the vicinity of the CDS start (**Figure 3C**). However, this pattern shifts towards a less biphasic profile with m^6^A becoming relatively more dominant around the stop codon in the later development stages when the methylation and expression levels increase (ks test, 5.87.10^−7^<*p*<5.35.10^−10^ depending on the cluster). Furthermore, these m^6^A-mRNAs have their methylation and expression dynamics strongly correlated (clusters 2, 3 and 4), whereas early DM transcripts without 5’-CDS methylation do not (cluster 1) (**Figure 3D**). This indicates that mRNA expression dynamics are correlated to m^6^A profile dynamics after, but not before, the zygotic genome activation. Together these results bring to light a regulation of gene expression by both the level and localization of m^6^A methylation across oyster developmental stages.

### Differentially m^6^A-methylated mRNAs bear the corresponding developmental functions

Gene ontology (GO) analyses of DM m^6^A-mRNAs shows an overall term enrichment related to developmental processes, like morphogenesis and mesoderm development (**Figure 4** and **Supplementary Data S1**). More specific functions correspond to each methylation cluster. The cluster 1 (cleavage) is enriched in terms related to the cell division whereas later clusters (gastrulation, tissue differentiation and metamorphosis) bear terms more reflecting signal transduction associated to cell differentiation such as the TGFβ/SMAD pathway. These results point to a functional implication of the m^6^A in the regulation of oyster developmental processes.

**Figure 4:**
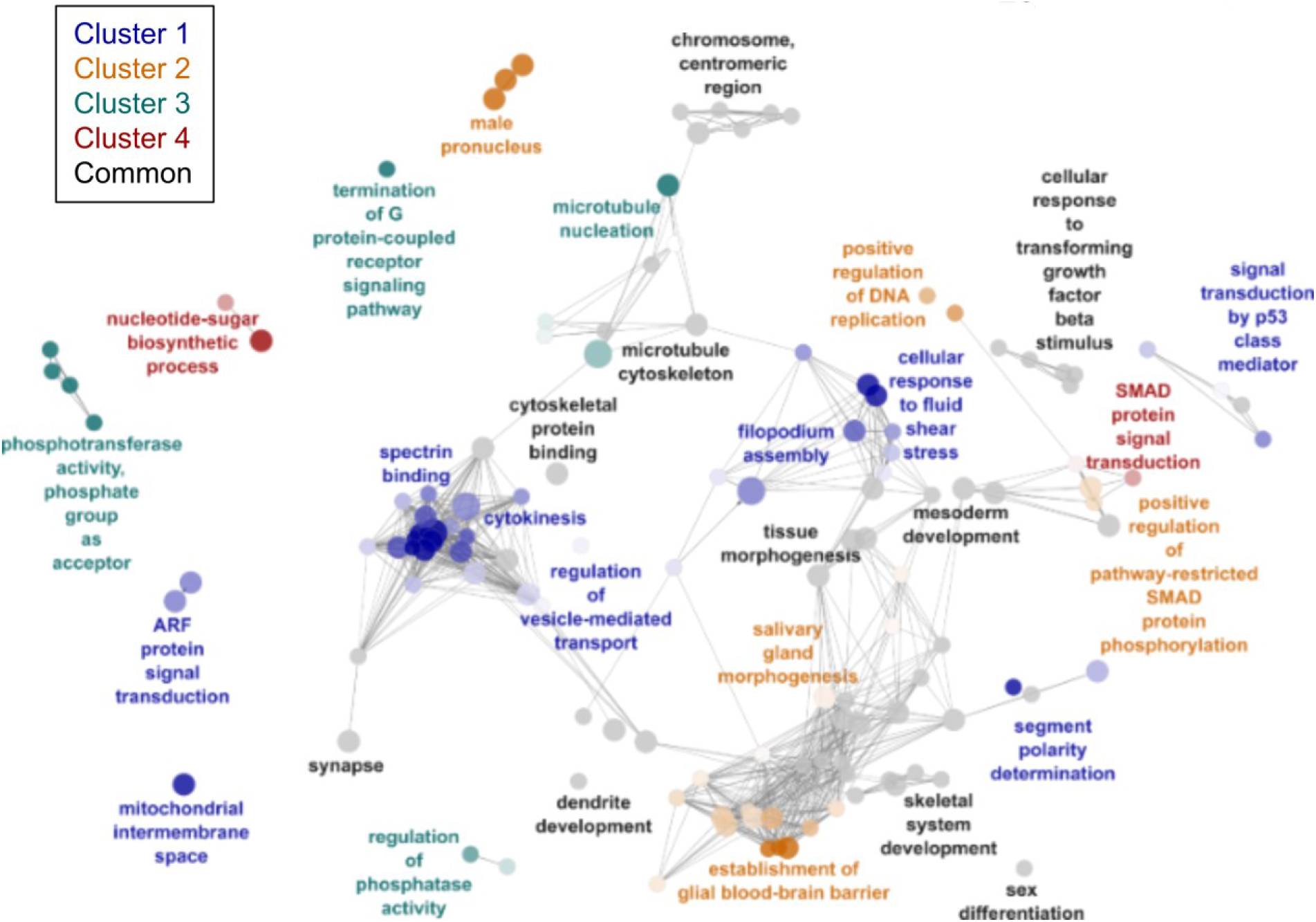
Functional annotation of differentially methylated mRNAs during oyster development. Enriched Gene Ontology terms (hypergeometric test, FDR<0.05) associated to differential m^6^A mRNA methylation across oyster development. Enriched terms corresponding to specific clusters are colored, common terms between clusters are black. Color intensity is relative to cluster specificity, with color limit set to 51% of genes inside the respective clusters. Circle diameter in inversely proportional to the *p*-value of the enrichment.

### Non-coding RNA transcript content is also associated to m^6^A-methylation profiles

Similar to mRNAs, lncRNA can be grouped into clusters according to the developmental chronology of their m^6^A methylation dynamics, with the highest methylation levels during the early stages for a cleavage cluster (cluster 1), during the gastrulation and tissue type differentiation (cluster 2), and at the pediveliger and spat stages for a metamorphosis cluster (cluster 3), respectively. A marked decrease of m^6^A levels at the morula stage is also observed for lncRNAs (**Figure 5A**). The methylation and expression levels are positively correlated overall (**Figure 5B**), although the methylation and expression dynamics are correlated only for the ‘late’ clusters 2 and 3, but not for the ‘early’ cluster 1 (**Figure 5C**).

**Figure 5:**
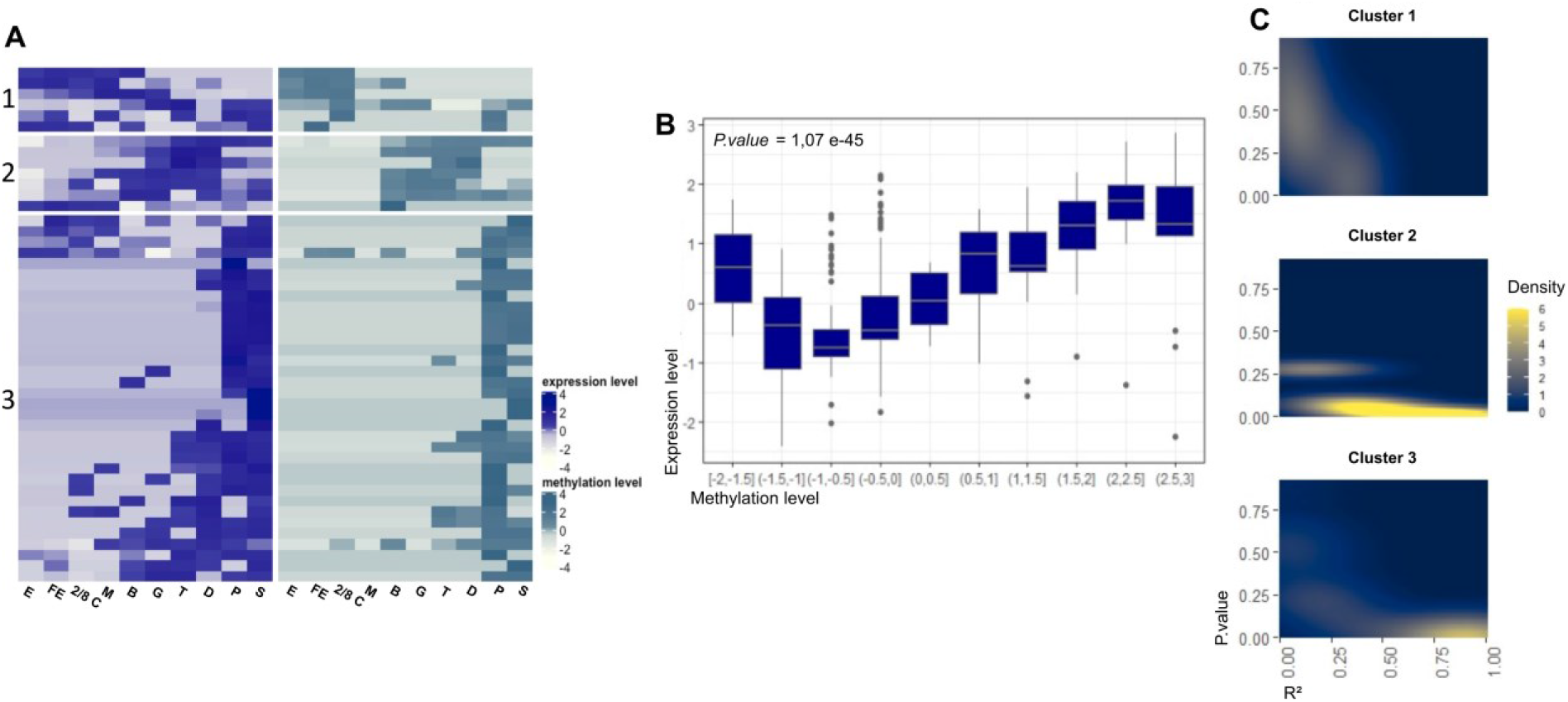
Dynamics of lncRNA expression and m^6^A methylation during oyster development. **A.** Normalized expression (blue) and m^6^A content (turquoise) of significantly methylated lncRNA genes during oyster development. Transcript variants were pooled for each gene. **B.** Correlation between expression (Y axis) and methylation (X axis) levels. The methylation level was divided into ten quantiles. **C**. Correlation plot of expression vs. methylation dynamics throughout development. The linear correlation of the methylation vs. expression variation was assessed for each gene across oyster development per cluster and the results are given as surface plots (R-square: X-axis; *p*-value: Y axis, density of genes: Z axis). E: Egg (oocyte), FE: Fertilized egg, 2/8C: Two to eight cell-stage, M: Morula, B: Blastula, G: Gastrula, T: Trochophore, D: D-larva, P: Pediveliger, S: Spat.

### Transposon RNA m^6^A-methylation is regulated across oyster development

The methylation of retrotransposon RNAs is mostly represented by the methylation of LTR and LINE groups (62.9% and 31.5% respectively). Their methylation levels display a peak at the blastula and trochophore stages for LTR and LINE respectively (**Figure 6A**). Both DNA and retrotransposon methylation levels were dynamic during oyster development (ANOVA *p*<0.05). TIR (37.1%), Helitron (32.5%) and Crypton (20.2%) are the three main DNA transposon types which transcripts are the most methylated. Their m^6^A levels are the highest around the gastrulation, and Helitron TEs exhibit an additional peak in pediveliger larvae (**Figure 6B**). These results show that the m^6^A-methylation of transposon transcripts display class-specific dynamics during oyster development that are different from mRNAs and lncRNAs.

**Figure 6:**
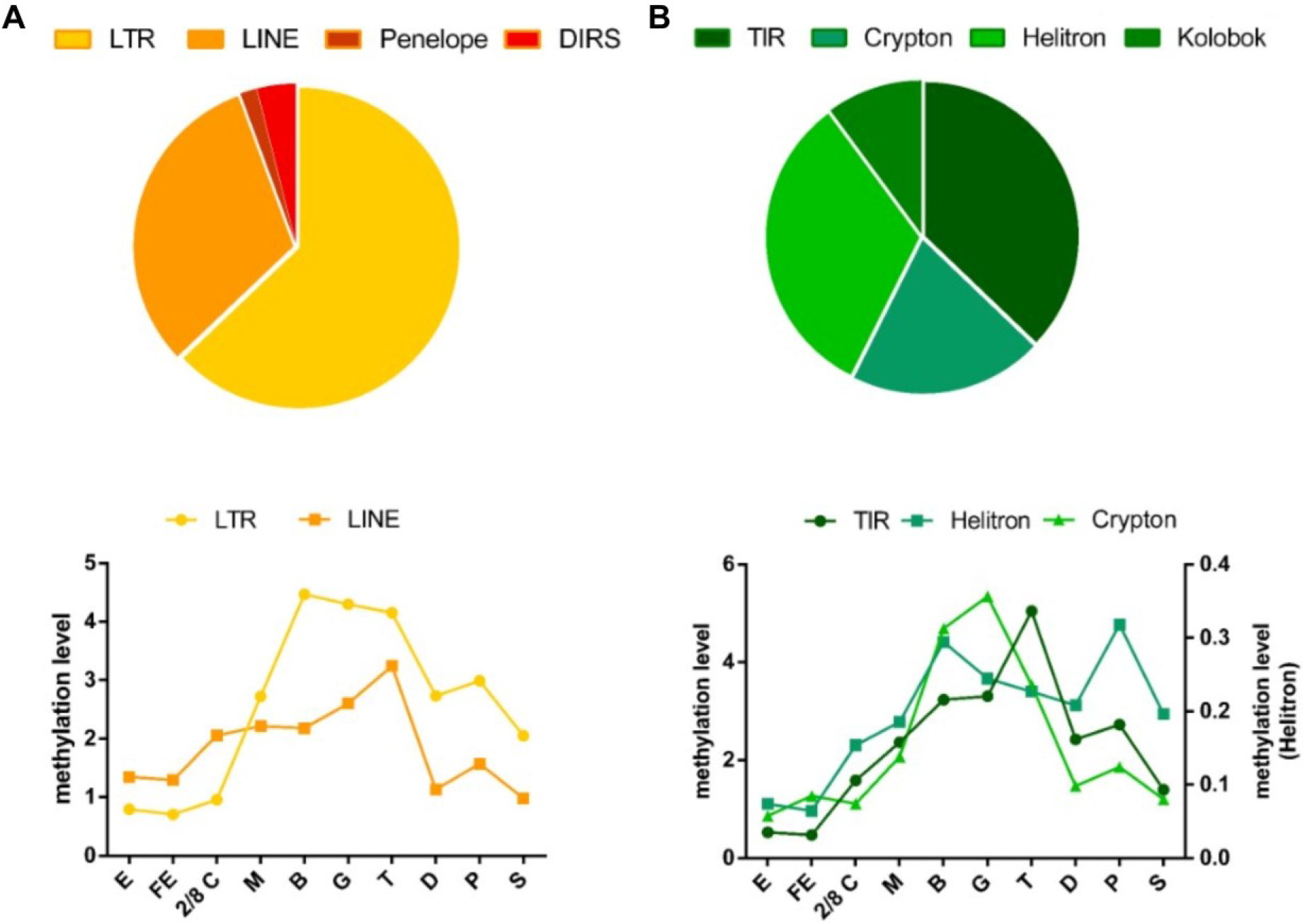
m^6^A dynamics of transposable element transcripts during oyster development. **A.** Relative representation of retrotranposon transcripts (top) and methylation level variation during oyster development (bottom). **B**. Relative representation of DNA TE transcripts (top) and methylation level variation during oyster development (bottom). Values are given as the mean of the two independent experiments. Only TEs representing more than 20% of their class are represented and error bars were omitted for clarity. Reference methylation level is the mean oocyte methylation level of Retro- and DNA transposons, respectively.

## Discussion

This work presents the profiling of m^6^A epitranscriptomes during the development of the oyster *Crassostrea gigas* from the egg to the completion of organogenesis. We demonstrate that m^6^A-methylation of RNA is dynamic throughout the oyster early life and specific to the RNA population considered. The methylation kinetics of mRNAs define clusters corresponding to the chronology of development, is correlated to gene expression level and dynamics, and the genes which transcripts display m^6^A-methylation regulation bear developmental functions. LncRNAs display similar profiles while transposable element transcripts have a specific signature with a peak in m^6^A levels during gastrulation. Our study provides evidence for a developmental significance of m^6^A-epitranscriptomes at various levels in the oyster.

The distinct oyster RNA classes present specific m^6^A signatures and vary in terms of methylation level and location. Indeed, while mRNAs are m^6^A-modified from the CDS start but mostly around the stop codon, lncRNAs do not present a biased methylation enrichment at their 3’ end. This result, suspected from our previous measurements of global m^6^A content in polyA vs. total RNAs (Le Franc et al. 2020), is more consistent with what is observed in human embryonic cell types (Xiao et al. 2019) than in insects where methylated adenosines are mostly lying in the 5’UTR and CDS (Wang et al. 2021; Kan et al. 2017; Li et al. 2019). This finding is rather surprising because insects are ecdysozoans which are considered the ‘sister clade’ of lophotrochozoans within protostomes. Nevertheless, this assumption is increasingly moderated by the closer resemablance of molecular features in mollusks and annelids to vertebrates, ie. deuterostomes, rather than to ecdysozoan features (Cho et al. 2010; Nederbragt et al. 2002; Saint-Carlier and Riviere 2015). Overall, RNA methyladenosines in the oyster are found within the motif ‘DRA*C’ which contains the GAC consensus sequence which is widely conserved across the evolution (Bodi et al. 2010; Luo et al. 2014; Lence et al. 2016; Wang et al. 2021; Meyer et al. 2012). The repetitive nature of TE sequences only allows less precise analysis of their m^6^A features compared to other RNA classes. However, we show that oyster DNA TE transcripts are significantly less methylated than RNA TEs, in line with observations in *Arabidopsis* (Wan et al. 2015).

Oyster epitranscriptomes are dynamic and associated to defined successive developmental steps, namely cleavage, gastrulation, tissue differentiation and metamorphosis. The similarity of m^6^A profiles in eggs before and after fertilization assesses poor contribution of sperm to embryonic m^6^A methylomes, consistent with the sperm scarce RNA content regarding the oocyte stock. Epitranscriptomes clearly shift during the developmental chronology, with a marked decrease of m^6^A levels in mRNA and lncRNA transcripts at the end of cleavage followed by a gradual increase, with a maximum methylation reached at the pre-metamorphosis stage that remains high afterwards in juveniles. The morula stage constitutes a pivot point where the methylation of mRNA and lncRNA, but not of TEs RNA, is strongly depleted. Interestingly, based on transcriptomic (Riviere et al. 2015) and 5mC-DNA methylation data (Riviere et al. 2017), this stage is considered the main window of maternal-to-zygotic transition (MZT) in the oyster, when most maternal RNA resources are consumed and early transcriptional events occur (McLean and Whiteley 1974). Interestingly, the marked epitranscriptomic features at cleavage and metamorphosis remind of 5mC-DNA dynamics (Riviere et al. 2017) and suggests interplay between epigenetic layers in the regulation of oyster development, possibly through competition for methyl donor (Shima et al. 2017) or histone-related pathways (Huang et al. 2019; Wang et al. 2018). This observation is reminiscent of the crosstalk between the 5mC-DNA status of the Pou5f1 pioneer stem cell factor promoter and the m^6^A methylation of its transcript in humans (Xiao et al. 2019), raising the question of the m^6^A-modified transcripts in the oyster development.

The m^6^A content and transcript abundance of DM mRNAs are positively correlated across oyster development, surprisingly contrasting with the situation in the honeybee where highly methylated transcripts exhibit downregulated expression (Wang et al. 2021). Overall, oyster DM mRNAs have functions related to developmental processes, further confirming the developmental significance of the epitranscriptomic regulation in the oyster. More precisely, DM mRNA transcripts group into 4 clusters that match the development steps described above, and which Gene Ontology annotations assess the functional relevance in the cognate developmental processes (cluster 1, cleavage; cluster 2, gastrulation; cluster 3, tissue differentiation; cluster 4, metamorphosis). However, these clusters do not always display similar methylation profiles neither correlation between m^6^A and expression level dynamics.

Cluster 1 mRNA transcripts are highly methylated and abundant during the cleavage. They are present in oocytes and therefore correspond to maternal transcripts. While their expression and methylation levels are dramatically reduced at the onset of MZT (morula stage), their m^6^A and transcript content dynamics are not significantly correlated, and they display predominant methylation around the stop codon throughout development. Such m^6^A profiles promote mRNA degradation *via* interaction with readers such as YTHDF proteins during cleavage in other species (Lee et al. 2020; Zhao et al. 2017), and we showed that an oyster YTHDF orthologue is present at high levels in this window (Le Franc et al. 2020). Besides, cluster 1 DM mRNAs bear functions in the cytoskeleton processes of cytokinesis. Together, these results indicate that the oyster MZT may be triggered by YTHDF-mediated decay of maternal mRNAs which translation is important for cell proliferation during cleavage. We hypothesize that the depletion of the cytokinesis machinery mRNA templates of maternal origin below a certain threshold, and/or the putative associated slowdown of cell divisions, may participate in triggering the zygotic genome activation. It remains to be deciphered whether and how such signals could be interpreted in terms of cell fate decisions.

The clusters 2, 3 and 4 respectively correspond to the gastrulation, the setting up of differentiated tissues and the metamorphosis. In contrast to cluster 1, their m^6^A pattern is dynamic and shifts from bimodal profiles in early stages to a late mostly unimodal m^6^A distribution around the stop codon. Such 5’-enriched m^6^A profiles are observed in plants (Luo et al. 2014), *Xenopus* (Sai et al. 2020), mice (Chang et al. 2017) and fruit flies (Worpenberg et al. 2021) and may be associated to increased RNA turnover and fast transcript processing depending on the positioning of methylation in the CDS or 5’UTR. Consistently, in developing oysters, the early bimodal m^6^A distribution is associated with a low transcript abundance. The 3’ patterns are associated to the recruitment of readers such as YTHDC1 (Kasowitz et al. 2018; Xu et al. 2014; Xiao et al. 2016), IGF2BP (Huang et al. 2018), or Prrc2a (Wu et al. 2018), and mediate translation initiation during gastrulation, as well as splicing and RNA trafficking during cell type differentiation. In the oyster, the later sequential shift of clusters towards more 3’-dominant methylation is correlated to an increased expression level of the DM mRNA transcripts. Furthermore, those transcripts that become successively methylated and expressed bear the cognate developmental functions such as mesoderm specification, morphogenesis, and signal transduction related to cell differentiation. In addition, readers including YTHDCs, IGF2BP, and Prrc2a are functional in the oyster and present in stage-specific sets after the MZT (Le Franc et al. 2020), and METTL3 expression is also dramatically increased after the MZT and sustained later on. Together, this strongly suggests that m^6^A regulates gene expression towards cell differentiation upon the zygotic genome activation, possibly by promoting translation *via* transcript selection and increased mRNA stability. As suggested by GO annotation, m^6^A likely participates in mesoderm formation during gastrulation *via* a METTL3-dependent regulation of epithelial to mesenchymal transition, as described in human cancer cells (Lin et al. 2019; Chen et al. 2017). Our findings are also consistent with the METTL3/YTHDF2-dependent osteoblast differentiation by SMAD inhibitor down-regulation in mammalian cells (Li et al. 2020; Zhang et al. 2020b). Regarding the conservation of the TGFβ pathway in oyster development (Herpin et al. 2005; Le Quéré et al. 2009), these results may also suggest feedback loops between TGFβ signaling and m^6^A regulation in the control of (pluri)potency during differentiation (Bertero et al. 2018).

LncRNAs methylation levels display similar developmental dynamics to mRNAs, although methyl-adenosines are not preferentially located at their 3’-end. Like for mRNAs, methylation levels in lncRNAs are also correlated to their transcript content and dynamics after the MZT. How lncRNA epitranscriptomes might participate in oyster development is less clear because oyster lncRNAs are poorly conserved and annotated. Besides, their limited number compared to mRNAs in addition to the limited resolution of MeRIP-seq did not allow the detection of m^6^A peak localization shifts. Nevertheless, the vertebrate writer complex cofactor RBM15/15B and the hnRNP2AB1 nuclear complex are conserved in the oyster (Le Franc et al. 2020). Therefore, we speculate that oyster lncRNA epitranscriptomes could regulate chromatin state and transcription in an m^6^A-dependent manner somewhat reminiscent of the *Xist*-mediated silencing (Patil et al. 2016; Coker et al. 2020), MALAT1-mediated stimulation (Brown et al. 2016; Liu et al. 2013) and/or chromosome-associated regulatory RNA (carRNA) switches in mammals (Liu et al. 2020). However, *Xist* is not conserved in the oyster and more work is required to test this hypothesis.

Transposon transcripts display class-specific dynamics during oyster development that are different from mRNAs and lncRNAs. Both RNA and DNA TEs which are poorly m^6^A modified during cleavage exhibit higher methylation during gastrulation and cell differentiation. Although the role of epitrancriptomes in transposon control remains largely unknown, this kinetics may suggest participation of m^6^A within the widely conserved RNA mechanisms of TE silencing required for cell differentiation (Shibata et al. 2016; Jankovics et al. 2018; Gerdes et al. 2016).

Altogether, our results demonstrate that oyster m^6^A-RNA epitranscriptomes are dynamic and RNA-class specific, and reveal their implication in oyster maternal-to-zygotic transition and sequential expression of genes required for gastrulation, cell differentiation and metamorphosis. Although additional studies would be required to determine the interplay of m^6^A-RNA with the epigenetic network and the underlying mechanisms, this first evidence of an epitranscriptomic regulation of development in a lophotrochozoan species allows a better understanding of developmental processes and their evolution.

## Materials & Methods

(see **supplemental methods** for detailed information)

### Animals

Broodstock oysters, embryos, larvae and spat were obtained at the IFREMER marine facility (Argenton, France) as previously described (Petton et al. 2015; Riviere et al. 2017). Briefly, gametes from conditioned mature broodstock specimen were obtained by gonad stripping and fertilization was triggered by the addition of ca.10 spermatozoids per oocyte. Embryos were transferred in oxygenated sterile sea water (SSW) tanks at 21 °C (<500 embryos/mL). The embryonic stages were determined by light microscopy observation: oocytes (E, immediately before sperm addition), fertilized oocytes (FE, immediately before transfer to tanks), two to eight cell embryos (2/8 C, ca. 1.5 hours post fertilization (hpf)), morula (M, ca. 4 hpf), blastula (B, ca. 6 hpf), gastrula (G, ca. 10 hpf), trochophore (T, ca 16 hpf) and D larvae (D, ca. 24 hpf), pediveliger (P, 20 days post-fertilization (dpf)) and spat (S, 25 dpf, post-metamorphosis) stages.

For each stage, 3 million embryos were collected and 1 million embryos (or 100 mg for P and S samples) were snap-frozen in liquid nitrogen directly of after resuspension in Tri-Reagent (Sigma-Aldrich, St Louis, MO, USA) (1 mL/10^6^ embryos). All samples were stored at −80 °C. Two distinct experiments were realized (February and March 2019) using the 126 and 140 broodstock animals, respectively.

### RNA extraction and fragmentation

RNA of each development stages was extracted using phenol-chloroform followed by affinity chromatography as previously described (Riviere et al. 2011). Briefly, embryos were ground in Tri-Reagent and RNA was purified using affinity chromatography (Nucleospin RNA Clean up kit, Macherey-Nagel, Duren, Germany). Potential contaminating DNA was removed by digestion with rDNase then RNA was purified using Beckman Coulter’s solid-phase reversible immobilization (SPRI) paramagnetic beads (Agencourt AMPure XP, Beckman Coulter, Brea, CA, USA) according to the manufacturer’s instructions.

The RNA (5 μg of total RNA from each stage) was heat-fragmented as previously described (Zeng et al. 2018) then purified using Beckman Coulter’s SPRI paramagnetic beads (Agencourt AMPure XP, Beckman Coulter). The fragment size (ca. 100 nucleotides) was verified by fluorescent capillary electrophoresis using an Agilent 2100 Bioanalyzer (Agilent RNA 6000 pico kit, Agilent, Santa Clara, CA, USA).

### MeRIP

The MeRIP was performed on fragmented RNA. Ten percent of the fragmented RNA solution (i.e. 500 ng of starting RNA before fragmentation) were not precipitated and used as the input fraction (Input), and 90% (i.e. 4.5 μg of starting RNA before fragmentation) were subjected to m6A-immunoprecipitation using m^6^A antibody-coupled (ABE572, Millipore, Burlington, MA, USA) protein A/G magnetic beads following the low/high salt procedure by Zeng and colleagues (Zeng et al. 2018). The immunoprecpitated RNA was purified using affinity chromatography (Nucleospin RNA Clean up kit, Macherey Nagel, France) and eluted with RNase free water. These immunoprecipitated RNAs correspond to the immunoprecipitated fraction (IP).

### RT-qPCR

To validate the immunoprecipitation of m^6^A methylated RNA, a RT-qPCR was performed on immunoprecipitated RNA pools composed of equal amounts of RNA from each developmental stage for the distinct experiments. Input and IP fractions were obtained as described above. The equivalent amount of 1.75 ng of Input and IP RNAs were used as starting template for the RT-qPCR protocol previously described (Riviere et al. 2011). Targets included one reference transcript (*ef1α*) and five conserved genes exhibiting distinct transcript m^6^A-methylation in vertebrates (*c-myc*, *klf*, *mettl3*, *hnrnpa2b1* and *oct4*). The methylation level was calculated as the normalized ratio of the IP and the Input signals by the formulae: 2^(-Δ(Ct IP-Ct Input)).

### Library preparation and sequencing

Amounts equivalent to 1.125 μg of starting RNA of the IP fraction and to the 50 ng of starting RNA of the Input fraction for each sample, respectively, were used for library construction using the SMARTer Stranded Total RNA-Seq Kit v.2 (634418, Pico Input Mammalian, Takara/Clontech, Japan) according to the manufacturer’s protocol without RNA fragmentation. The ribosomal cDNA depletion step and a final cDNA amplification of 16 cycles were performed. Paired-end 150-bp sequencing of Input and IP cDNA libraries of each sample were conducted on an Illumina HiSeq 4000 platform at the Genome Quebec innovation Center (McGill University, Montréal, Canada).

### MeRIP-seq Data analyses

Read quality was evaluated using FastQC (v.0.11.7) and MultiQC (v.1.7) and adaptor sequences and low-quality reads were removed using Trimmommatic (v.0.38). The remaining reads were aligned to the oyster genome v.9 (GCF_000297895.1) and uniquely mapped reads were counted using STAR (v.2.7.3a; -quantMode GeneCounts) (Dobin et al. 2013). Messenger and non-coding RNA were identified from the gene annotation and TEs were identified using the RepeatMasker annotation output provided with assembly data. The expression level in all Input samples was expressed in TPM (Transcripts per Million) (Li et al. 2010). The m^6^A-enriched peaks were identified on uniquely mapped reads of the IP samples using Samtools (v.1.9) and MeTPeak R package (Cui et al. 2016a) (FDR < 5%) with the paired Input samples as controls. The methylation level of m^6^A-enriched RNAs corresponds to the IP/Input fold-change provided by MeTPeak. The distribution of m^6^A across mRNAs and ncRNAs was visualized using Guitar plots (Cui et al. 2016b). The methylation level of TEs was assessed as the ratio of IP/Input with reads per gene expressed in TPM. Transcript variants were pooled for each gene and only transcripts present in the two distinct development experiments were considered expressed or methylated in Input and IP data, respectively. The m^6^A motif was searched using Homer (v.4.10.4). All peaks mapped on mRNAs and ncRNAs were used as the target sequences and the background sequences were constituted of 5% of the Input reads using Samtools (v.1.9).

### Gene ontology analysis

The RNA sequences identified as differentially methylated across oyster development were identified using BlastN (Altschul et al. 1997) against the GigaTON reference transcriptome database (Riviere et al. 2015) with default settings. Gene ontology (GO) analyses were carried out with the GO annotations obtained from the GigaTON database gene universe (Riviere et al. 2015). GO-term enrichment tests were performed using ClueGO plugin (v.2.5.7) (Bindea et al. 2009) on Cytoscape (v.3.8.0). The hypergeometric test with a FDR <5 % was used to consider significant GO term enrichment.

### Statistical analyses and graph production

The m^6^A enrichment and the variation of the methylation level across oyster development were analyzed using one-way ANOVA (factor: development stage) followed by Bonferroni’s post-hoc test. Methylation profiles were compared using Kolmogorov-Smirnoff tests. Correlations were estimated using linear regression. A *p*-value <0.05 was considered significant. Values were log-centered reduced for heatmap production. Statistical analyses were performed using R (v.3.6.3) and RStudio (v.1.0.153) softwares. The R packages *eulerr* (Larsson 2019), *Complexheatmap* (Gu et al. 2016), *ggplot2* (Gómez-Rubio 2017), *Guitar* (Cui et al. 2016b), *PCATools* (Blighe and Lewis 2019), *corrplot* (Wei and Simko 2017) and Graphpad v.6 (Prism) software were used for figure production.

## Supporting information

Expanded methods

## Abbreviations

(m^6^A): *N*^6^-methyladenosine
(TE): Transposable elements
(UTR): Untranslated transcribed region
(CDS): Coding sequences
(mRNA): messenger RNA
(lncRNA): long non-coding RNA
(misc. RNA): miscellaneous RNA
(tRNA): transfer RNA
(SSW): Sterile sea water
(egg, E): Oocyte
(FE): Fertilized oocyte
(2/8 C): Two to eight cell embryo
(M): Morula
(B): Blastula
(G): Gastrula
(T): Trochophore
(D): D larvae
(P): Pedeveliger
(S): Spat
(hpf): Hours post fertilization
(dpf): Days post fertilization
(GO): Gene ontology
(MZT): Maternal-to-zygotic transition
(DM): Differentially methylated
(Input): Input fraction
(IP): Immunoprecipitated fraction
(TPM): Transcripts per million

## Data availability

All data relative to this study are publicly available at the NCBI Gene Expression Omnibus under the accession number GSE180388.

## Competing interest statement

The authors declare they have no competing interest.

## Aknowledgements

This work was supported by the French national program CNRS EC2CO ECODYN (Ecosphère Continentale et Côtière ‘HERITAGe’ to G. Rivière) and the Council of the Normandy Region (RIN ECUME to P. Favrel).

## Author contributions

GR and LLF designed the experiment and analyzed the data. LLF, GR and BP performed sampling and benchwork. LLF, GR, PF and BP wrote and edited the manuscript.

## Notes

### Competing Interest Statement

The authors have declared no competing interest.

