## Supplemental figure 1 for "Profiling of *N*^6^-methyladenosine dynamics indicates regulation of oyster development by m^6^A-RNA epitranscriptomes"

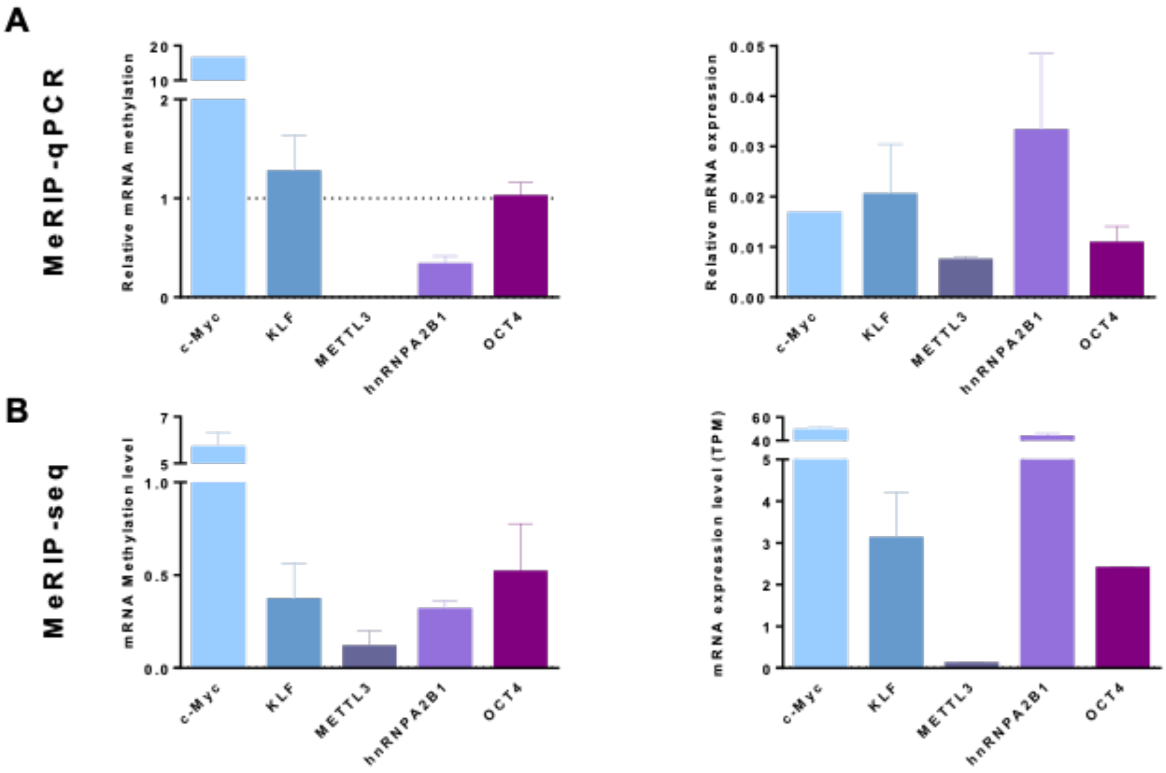


**Supplemental_Figure_S1: Validation of m$^6$A immunoprecipitation transcripts**
**A**. Relative methylation (left) and expression level (right) of targeted transcripts c-Myc, KLF, METTL3, hnRNPA2B1 and Oct4 quantified by MeRIP-qPCR (left) and qPCR (right) on a pool of oyster development. The dotted line represents the minimal value to consider an enrichment of m$^6$A-RNA. **B**. Methylation (left) and expression level in TPM (right) of targeted transcripts c-Myc, KLF, METTL3, hnRNPA2B1 and Oct4 quantified by MeRIP-seq (left) and RNA-seq (left) on a pool of oyster development. Error bars represent the standard deviation.
