## Supplemental Table 1 for "Profiling of *N*^6^-methyladenosine dynamics indicates regulation of oyster development by m^6^A-RNA epitranscriptomes"

**Supplemental_Table_S1: Complete list of significant GO terms in each cluster for m6A-mRNAs (FDR < 0,05)**

| Cluster | GOID | GOTerm | Ontology Source | FDR | % Associated Genes | Nr. Genes |
|---|---|---|---|---|---|---|
| None Specific Cluster | GO:0007548 | sex differentiation | BP | 0,02 | 11,78 | 41,00 |
| None Specific Cluster | GO:0008092 | cytoskeletal protein binding | MF | 0,00 | 14,29 | 35,00 |
| None Specific Cluster | GO:0016358 | dendrite development | BP | 0,04 | 14,42 | 15,00 |
| None Specific Cluster | GO:0060627 | regulation of vesicle-mediated transport | BP | 0,02 | 18,33 | 11,00 |
| None Specific Cluster | GO:0097193 | intrinsic apoptotic signaling pathway | BP | 0,05 | 18,60 | 8,00 |
| None Specific Cluster | GO:0008630 | intrinsic apoptotic signaling pathway in response to DNA damage | BP | 0,03 | 26,09 | 6,00 |
| None Specific Cluster | GO:0098858 | actin-based cell projection | CC | 0,01 | 22,92 | 11,00 |
| None Specific Cluster | GO:0007391 | dorsal closure | BP | 0,01 | 37,50 | 6,00 |
| None Specific Cluster | GO:0120031 | plasma membrane bounded cell projection assembly | BP | 0,01 | 16,98 | 18,00 |
| None Specific Cluster | GO:0051489 | regulation of filopodium assembly | BP | 0,01 | 30,43 | 7,00 |
| None Specific Cluster | GO:0071363 | cellular response to growth factor stimulus | BP | 0,03 | 11,38 | 43,00 |
| None Specific Cluster | GO:0071560 | cellular response to transforming growth factor beta stimulus | BP | 0,01 | 12,89 | 33,00 |
| None Specific Cluster | GO:0007178 | transmembrane receptor protein serine/threonine kinase signaling pathway | BP | 0,04 | 11,66 | 33,00 |
| None Specific Cluster | GO:0007179 | transforming growth factor beta receptor signaling pathway | BP | 0,02 | 12,40 | 31,00 |
| None Specific Cluster | GO:0045944 | positive regulation of transcription by RNA polymerase II | BP | 0,02 | 11,29 | 49,00 |
| None Specific Cluster | GO:0000775 | chromosome, centromeric region | CC | 0,00 | 15,23 | 37,00 |
| None Specific Cluster | GO:0000792 | heterochromatin | CC | 0,01 | 13,92 | 27,00 |
| None Specific Cluster | GO:0000793 | condensed chromosome | CC | 0,03 | 12,24 | 30,00 |
| None Specific Cluster | GO:0005819 | spindle | CC | 0,03 | 14,18 | 19,00 |
| None Specific Cluster | GO:0045815 | positive regulation of gene expression, epigenetic | BP | 0,05 | 12,72 | 22,00 |
| None Specific Cluster | GO:0005721 | pericentric heterochromatin | CC | 0,01 | 14,12 | 25,00 |
| None Specific Cluster | GO:0034401 | chromatin organization involved in regulation of transcription | BP | 0,05 | 12,50 | 23,00 |
| None Specific Cluster | GO:0001501 | skeletal system development | BP | 0,01 | 14,29 | 26,00 |
| None Specific Cluster | GO:0051216 | cartilage development | BP | 0,01 | 16,96 | 19,00 |
| None Specific Cluster | GO:0072657 | protein localization to membrane | BP | 0,03 | 15,46 | 15,00 |

| | | | | | | |
|---|---|---|---|---|---|---|
| None Specific Cluster | GO:0002062 | chondrocyte differentiation | BP | 0,01 | 18,48 | 17,00 |
| None Specific Cluster | GO:0032330 | regulation of chondrocyte differentiation | BP | 0,00 | 20,73 | 17,00 |
| None Specific Cluster | GO:0006605 | protein targeting | BP | 0,00 | 16,20 | 23,00 |
| None Specific Cluster | GO:0006612 | protein targeting to membrane | BP | 0,00 | 23,53 | 12,00 |
| None Specific Cluster | GO:0006607 | NLS-bearing protein import into nucleus | BP | 0,01 | 19,74 | 15,00 |
| None Specific Cluster | GO:0007389 | pattern specification process | BP | 0,02 | 13,07 | 26,00 |
| None Specific Cluster | GO:0048736 | appendage development | BP | 0,03 | 15,56 | 14,00 |
| None Specific Cluster | GO:0007498 | mesoderm development | BP | 0,00 | 28,89 | 13,00 |
| None Specific Cluster | GO:0048332 | mesoderm morphogenesis | BP | 0,00 | 33,33 | 8,00 |
| None Specific Cluster | GO:0045669 | positive regulation of osteoblast differentiation | BP | 0,00 | 46,15 | 6,00 |
| None Specific Cluster | GO:0090100 | positive regulation of transmembrane receptor protein serine/threonine kinase signaling pathway | BP | 0,03 | 23,33 | 7,00 |
| None Specific Cluster | GO:0032507 | maintenance of protein location in cell | BP | 0,00 | 20,90 | 14,00 |
| None Specific Cluster | GO:0000775 | chromosome, centromeric region | CC | 0,00 | 15,23 | 37,00 |
| None Specific Cluster | GO:0005819 | spindle | CC | 0,03 | 14,18 | 19,00 |
| None Specific Cluster | GO:0015630 | microtubule cytoskeleton | CC | 0,01 | 12,28 | 48,00 |
| None Specific Cluster | GO:0099513 | polymeric cytoskeletal fiber | CC | 0,02 | 13,21 | 28,00 |
| None Specific Cluster | GO:0005874 | microtubule | CC | 0,01 | 14,20 | 25,00 |
| None Specific Cluster | GO:0051258 | protein polymerization | BP | 0,00 | 18,02 | 20,00 |
| None Specific Cluster | GO:0045202 | synapse | CC | 0,00 | 14,59 | 34,00 |
| None Specific Cluster | GO:0005911 | cell-cell junction | CC | 0,00 | 14,94 | 39,00 |
| None Specific Cluster | GO:0032507 | maintenance of protein location in cell | BP | 0,00 | 20,90 | 14,00 |
| None Specific Cluster | GO:0044304 | main axon | CC | 0,00 | 21,21 | 14,00 |
| None Specific Cluster | GO:0045211 | postsynaptic membrane | CC | 0,01 | 17,65 | 18,00 |
| None Specific Cluster | GO:0072657 | protein localization to membrane | BP | 0,03 | 15,46 | 15,00 |
| None Specific Cluster | GO:0006605 | protein targeting | BP | 0,00 | 16,20 | 23,00 |
| None Specific Cluster | GO:0006612 | protein targeting to membrane | BP | 0,00 | 23,53 | 12,00 |
| None Specific Cluster | GO:0007389 | pattern specification process | BP | 0,02 | 13,07 | 26,00 |

| | | | | | | |
|---|---|---|---|---|---|---|
| None Specific Cluster | GO:0009791 | post-embryonic development | BP | 0,00 | 21,92 | 16,00 |
| None Specific Cluster | GO:0007552 | metamorphosis | BP | 0,00 | 26,83 | 11,00 |
| None Specific Cluster | GO:0048729 | tissue morphogenesis | BP | 0,00 | 14,89 | 42,00 |
| None Specific Cluster | GO:0048736 | appendage development | BP | 0,03 | 15,56 | 14,00 |
| None Specific Cluster | GO:0002165 | instar larval or pupal development | BP | 0,00 | 28,57 | 12,00 |
| None Specific Cluster | GO:0007498 | mesoderm development | BP | 0,00 | 28,89 | 13,00 |
| None Specific Cluster | GO:0001655 | urogenital system development | BP | 0,03 | 13,73 | 21,00 |
| None Specific Cluster | GO:0002009 | morphogenesis of an epithelium | BP | 0,00 | 15,56 | 40,00 |
| None Specific Cluster | GO:0060541 | respiratory system development | BP | 0,00 | 18,10 | 19,00 |
| None Specific Cluster | GO:0007444 | imaginal disc development | BP | 0,01 | 22,73 | 10,00 |
| None Specific Cluster | GO:0016331 | morphogenesis of embryonic epithelium | BP | 0,02 | 18,64 | 11,00 |
| None Specific Cluster | GO:0043296 | apical junction complex | CC | 0,01 | 24,32 | 9,00 |
| None Specific Cluster | GO:0048732 | gland development | BP | 0,01 | 13,76 | 26,00 |
| None Specific Cluster | GO:0060562 | epithelial tube morphogenesis | BP | 0,00 | 17,39 | 24,00 |
| None Specific Cluster | GO:0007424 | open tracheal system development | BP | 0,02 | 21,62 | 8,00 |
| None Specific Cluster | GO:0022612 | gland morphogenesis | BP | 0,01 | 17,12 | 19,00 |
| None Specific Cluster | GO:0030324 | lung development | BP | 0,03 | 17,86 | 10,00 |
| None Specific Cluster | GO:0007043 | cell-cell junction assembly | BP | 0,03 | 15,91 | 14,00 |
| None Specific Cluster | GO:0007391 | dorsal closure | BP | 0,01 | 37,50 | 6,00 |
| None Specific Cluster | GO:0007560 | imaginal disc morphogenesis | BP | 0,01 | 24,24 | 8,00 |
| None Specific Cluster | GO:0120031 | plasma membrane bounded cell projection assembly | BP | 0,01 | 16,98 | 18,00 |
| None Specific Cluster | GO:0007435 | salivary gland morphogenesis | BP | 0,00 | 34,62 | 9,00 |
| None Specific Cluster | GO:0061333 | renal tubule morphogenesis | BP | 0,00 | 34,48 | 10,00 |
| None Specific Cluster | GO:0072073 | kidney epithelium development | BP | 0,03 | 19,15 | 9,00 |
| None Specific Cluster | GO:0051258 | protein polymerization | BP | 0,00 | 18,02 | 20,00 |
| None Specific Cluster | GO:0051489 | regulation of filopodium assembly | BP | 0,01 | 30,43 | 7,00 |
| None Specific Cluster | GO:0007476 | imaginal disc-derived wing morphogenesis | BP | 0,02 | 25,00 | 7,00 |
| Specific for Cluster #1 | GO:0005758 | mitochondrial intermembrane space | CC | 0,00 | 50,00 | 5,00 |

| | | | | | | |
|---|---|---|---|---|---|---|
| Specific for Cluster #1 | GO:0007367 | segment polarity determination | BP | 0,03 | 50,00 | 3,00 |
| Specific for Cluster #1 | GO:0032011 | ARF protein signal transduction | BP | 0,00 | 47,06 | 8,00 |
| Specific for Cluster #1 | GO:0032012 | regulation of ARF protein signal transduction | BP | 0,00 | 47,06 | 8,00 |
| Specific for Cluster #1 | GO:0072331 | signal transduction by p53 class mediator | BP | 0,01 | 25,81 | 8,00 |
| Specific for Cluster #1 | GO:0031571 | mitotic G1 DNA damage checkpoint | BP | 0,05 | 30,77 | 4,00 |
| Specific for Cluster #1 | GO:0046847 | filopodium assembly | BP | 0,00 | 34,21 | 13,00 |
| Specific for Cluster #1 | GO:0030507 | spectrin binding | MF | 0,00 | 58,33 | 7,00 |
| Specific for Cluster #1 | GO:0072659 | protein localization to plasma membrane | BP | 0,00 | 27,78 | 10,00 |
| Specific for Cluster #1 | GO:0008154 | actin polymerization or depolymerization | BP | 0,02 | 17,91 | 12,00 |
| Specific for Cluster #1 | GO:0000910 | cytokinesis | BP | 0,01 | 19,67 | 12,00 |
| Specific for Cluster #1 | GO:0030315 | T-tubule | CC | 0,00 | 43,75 | 7,00 |
| Specific for Cluster #1 | GO:0030507 | spectrin binding | MF | 0,00 | 58,33 | 7,00 |
| Specific for Cluster #1 | GO:0001508 | action potential | BP | 0,05 | 15,94 | 11,00 |
| Specific for Cluster #1 | GO:0033268 | node of Ranvier | CC | 0,03 | 17,86 | 10,00 |
| Specific for Cluster #1 | GO:0042383 | sarcolemma | CC | 0,01 | 21,74 | 10,00 |
| Specific for Cluster #1 | GO:0045838 | positive regulation of membrane potential | BP | 0,00 | 42,86 | 6,00 |
| Specific for Cluster #1 | GO:0000281 | mitotic cytokinesis | BP | 0,00 | 61,54 | 8,00 |
| Specific for Cluster #1 | GO:0006892 | post-Golgi vesicle-mediated transport | BP | 0,03 | 19,57 | 9,00 |
| Specific for Cluster #1 | GO:0071709 | membrane assembly | BP | 0,01 | 30,00 | 6,00 |
| Specific for Cluster #1 | GO:0086001 | cardiac muscle cell action potential | BP | 0,00 | 43,75 | 7,00 |
| Specific for Cluster #1 | GO:0014704 | intercalated disc | CC | 0,01 | 30,00 | 6,00 |
| Specific for Cluster #1 | GO:0032414 | positive regulation of ion transmembrane transporter activity | BP | 0,02 | 25,93 | 7,00 |
| Specific for Cluster #1 | GO:0072659 | protein localization to plasma membrane | BP | 0,00 | 27,78 | 10,00 |
| Specific for Cluster #1 | GO:0072660 | maintenance of protein location in plasma membrane | BP | 0,00 | 66,67 | 6,00 |
| Specific for Cluster #1 | GO:1900827 | positive regulation of membrane depolarization during cardiac muscle cell action potential | BP | 0,00 | 85,71 | 6,00 |
| Specific for Cluster #1 | GO:0043001 | Golgi to plasma membrane protein transport | BP | 0,00 | 54,55 | 6,00 |
| Specific for Cluster #1 | GO:0006813 | potassium ion transport | BP | 0,03 | 17,86 | 10,00 |

| | | | | | | |
|---|---|---|---|---|---|---|
| Specific for Cluster #1 | GO:0071498 | cellular response to fluid shear stress | BP | 0,00 | 83,33 | 5,00 |
| Specific for Cluster #1 | GO:0003785 | actin monomer binding | MF | 0,00 | 71,43 | 5,00 |
| Specific for Cluster #1 | GO:0048286 | lung alveolus development | BP | 0,00 | 42,86 | 6,00 |
| Specific for Cluster #1 | GO:0061005 | cell differentiation involved in kidney development | BP | 0,01 | 31,82 | 7,00 |
| Specific for Cluster #1 | GO:0008154 | actin polymerization or depolymerization | BP | 0,02 | 17,91 | 12,00 |
| Specific for Cluster #1 | GO:2001013 | epithelial cell proliferation involved in renal tubule morphogenesis | BP | 0,00 | 66,67 | 6,00 |
| Specific for Cluster #1 | GO:0046847 | filopodium assembly | BP | 0,00 | 34,21 | 13,00 |
| Specific for Cluster #1 | GO:0072080 | nephron tubule development | BP | 0,02 | 22,22 | 8,00 |
| Specific for Cluster #2 | GO:0045740 | positive regulation of DNA replication | BP | 0,03 | 33,33 | 4,00 |
| Specific for Cluster #2 | GO:0010369 | chromocenter | CC | 0,00 | 80,00 | 4,00 |
| Specific for Cluster #2 | GO:0001939 | female pronucleus | CC | 0,00 | 66,67 | 4,00 |
| Specific for Cluster #2 | GO:0001940 | male pronucleus | CC | 0,00 | 80,00 | 4,00 |
| Specific for Cluster #2 | GO:0060389 | pathway-restricted SMAD protein phosphorylation | BP | 0,01 | 38,46 | 5,00 |
| Specific for Cluster #2 | GO:0010862 | positive regulation of pathway-restricted SMAD protein phosphorylation | BP | 0,00 | 62,50 | 5,00 |
| Specific for Cluster #2 | GO:0018210 | peptidyl-threonine modification | BP | 0,01 | 33,33 | 6,00 |
| Specific for Cluster #2 | GO:0007527 | adult somatic muscle development | BP | 0,00 | 100,00 | 3,00 |
| Specific for Cluster #2 | GO:0009954 | proximal/distal pattern formation | BP | 0,01 | 38,46 | 5,00 |
| Specific for Cluster #2 | GO:0070160 | tight junction | CC | 0,01 | 24,24 | 8,00 |
| Specific for Cluster #2 | GO:0008362 | chitin-based embryonic cuticle biosynthetic process | BP | 0,03 | 50,00 | 3,00 |
| Specific for Cluster #2 | GO:0005918 | septate junction | CC | 0,00 | 62,50 | 5,00 |
| Specific for Cluster #2 | GO:0035151 | regulation of tube size, open tracheal system | BP | 0,01 | 35,71 | 5,00 |
| Specific for Cluster #2 | GO:0005920 | smooth septate junction | CC | 0,01 | 75,00 | 3,00 |
| Specific for Cluster #2 | GO:0043297 | apical junction assembly | BP | 0,00 | 42,86 | 6,00 |
| Specific for Cluster #2 | GO:0120192 | tight junction assembly | BP | 0,00 | 42,86 | 6,00 |
| Specific for Cluster #2 | GO:0060857 | establishment of glial blood-brain barrier | BP | 0,00 | 80,00 | 4,00 |
| Specific for Cluster #2 | GO:0035317 | imaginal disc-derived wing hair organization | BP | 0,05 | 30,77 | 4,00 |
| Specific for Cluster #3 | GO:0038032 | termination of G protein-coupled receptor signaling pathway | BP | 0,04 | 42,86 | 3,00 |
| Specific for Cluster #3 | GO:0010921 | regulation of phosphatase activity | BP | 0,02 | 22,22 | 8,00 |

| | | | | | | |
|---|---|---|---|---|---|---|
| Specific for Cluster #3 | GO:0010923 | negative regulation of phosphatase activity | BP | 0,02 | 27,27 | 6,00 |
| Specific for Cluster #3 | GO:0016776 | phosphotransferase activity, phosphate group as acceptor | MF | 0,02 | 33,33 | 5,00 |
| Specific for Cluster #3 | GO:0004550 | nucleoside diphosphate kinase activity | BP | 0,03 | 50,00 | 3,00 |
| Specific for Cluster #3 | GO:0006183 | GTP biosynthetic process | BP | 0,03 | 50,00 | 3,00 |
| Specific for Cluster #3 | GO:0006228 | UTP biosynthetic process | BP | 0,03 | 50,00 | 3,00 |
| Specific for Cluster #3 | GO:0031109 | microtubule polymerization or depolymerization | BP | 0,01 | 27,59 | 8,00 |
| Specific for Cluster #3 | GO:0034453 | microtubule anchoring | BP | 0,00 | 61,54 | 8,00 |
| Specific for Cluster #3 | GO:0007020 | microtubule nucleation | BP | 0,00 | 60,00 | 6,00 |
| Specific for Cluster #3 | GO:0034453 | microtubule anchoring | BP | 0,00 | 61,54 | 8,00 |
| Specific for Cluster #4 | GO:0004360 | glutamine-fructose-6-phosphate transaminase (isomerizing) activity | MF | 0,00 | 100,00 | 3,00 |
| Specific for Cluster #4 | GO:0009226 | nucleotide-sugar biosynthetic process | BP | 0,01 | 44,44 | 4,00 |
| Specific for Cluster #4 | GO:0060395 | SMAD protein signal transduction | BP | 0,01 | 57,14 | 4,00 |
