## Supplementary material for "Profiling of *N*^6^-methyladenosine dynamics indicates regulation of oyster development by m^6^A-RNA epitranscriptomes": Expanded methods

### Supplemental Methods

#### Animals:

Broodstock oysters, embryos, larvae and spat were obtained at the IFREMER marine facility (Argenton, France) as previously described (Petton et al. 2015; Riviere et al. 2017). Briefly, gametes of mature broodstock oysters were obtained by stripping the gonads and filtering the recovered material on a 60 µm mesh to remove large debris. Oocytes were collected as the remaining fraction on a 20 µm mesh and spermatozoa as the passing fraction on a 20 µm mesh. Oocytes were pre-incubated in 5 L of UV-treated and 1 µm filtered sterile sea water (SSW) at 21 °C until germinal vesicle breakdown. Fertilization was triggered by the addition of ca.10 spermatozoids per oocyte. After the expulsion of the second polar body was assessed by light microscopy, embryos were transferred in 150 L tanks of oxygenated SSW at 21 °C. The embryonic stages were determined by light microscopy observation. The embryonic stages collected were oocytes (E, immediately before sperm addition), fertilized oocytes (FE, immediately before transfer to 150 L tanks), two to eight cell embryos (2/8 C, ca. 1.5 hours post fertilization (hpf)), morula (M, ca. 4 hpf), blastula (B, ca. 6 hpf), gastrula (G, ca. 10 hpf), trochophore (T, ca 16 hpf) and D larvae (D, ca. 24 hpf). For pediveliger (P) and spat (S) stages, D-larvae were collected and reared in a flow-through rearing system at 25 °C in SSW. At the end of the pelagic stage, competent larvae were collected on a 100 µm to allow the larval settlement. At 20 day post fertilization (dpf) the pediveliger stage was sampled as the remaining fraction on a 400 µm mesh. The post-larvae were maintained in a downwelling system. Then, at 25 dpf the spat stage is sampled after metamorphosis as the remaining fraction on a 400 µm mesh.

For each embryonic stage, 3 million embryos were collected as the remaining fraction on a 20 µm mesh and centrifuged at 123 g for 5 min at room temperature. Supernatant was discarded and samples of 1 million embryos were then snap-frozen in liquid nitrogen directly of after resuspension in Tri-Reagent (Sigma-Aldrich, St Louis, MO, USA) (1 mL/$10^6$ embryos). For pediveliger and spat samples, 100 mg of each stage were resuspended in Tri-Reagent (Sigma-Aldrich) (1 mL/100 mg) and snap-frozen in liquid nitrogen. All samples were stored at -80 °C. Two distinct experiments were realized (February and March 2019) using the gametes of 126 and 140 broodstock animals, respectively.

**RNA extraction and fragmentation:**

RNA of each development stages was extracted using phenol-chloroform followed by affinity chromatography as previously described (Riviere et al. 2011). Briefly, embryos were ground in Tri-Reagent and RNA was purified using affinity chromatography (Nucleospin RNA Clean up kit, Macherey-Nagel, Duren, Germany). Potential contaminating DNA was removed by digestion with rDNase (Macherey-Nagel) according to the manufacturer’s instructions for 15 min at 37 °C then RNA was purified using Beckman Coulter’s solid-phase reversible immobilization (SPRI) paramagnetic beads (AgencourtAMPure XP, Beckman Coulter, Brea, CA, USA) according to the manufacturer’s instructions. Briefly, paramagnetic beads and RNAs were mixed slowly and incubated 5 min at room temperature followed by 2 min on a magnetic rack. Cleared supernatant was removed, and beads were washed three times with 70 % ethanol. After 4 min of drying at room temperature, RNAs were mixed slowly with RNase free water and incubated for 1 min at room temperature on the magnetic rack. Eluted total RNA was stored at -80 °C.

The RNA was heat-fragmented as previously described (Zeng et al. 2018). Briefly, for each RNA sample (development stages and pools), 5 µg of total RNA was suspended in 18 µl of RNase free water. The RNA was fragmented by the addition of 2 µl of fragmentation buffer (100 mM Tris-HCl pH 7.0, 100 mM ZnCl2, DEPC water) and incubation for 2 min at 70 °C. After the incubation, 2 µl of EDTA 0.5 M was immediately added and RNAs were incubated on ice for 2 min to stop the reaction. Then, RNAs were purified using Beckman Coulter's SPRI paramagnetic beads (Agencourt AMPure XP, Beckman Coulter) as previously described and eluted in 50 µL of RNAse-free water. The fragment size (ca. 100 nucleotides) was verified by fluorescent capillary electrophoresis using an Agilent 2100 Bioanalyzer (Agilent RNA 6000 pico kit, Agilent, Santa Clara, CA, USA) according to the manufacturer's instruction.

**MeRIP:**

The MeRIP-seq of total RNA was performed on fragmented RNA. Of the 50 µL fragmented RNA solution, 5 µl (i.e. corresponding to 500 ng of starting RNA before fragmentation) was non immunoprecipitated and used as the input fraction (Input), and 44 µL (i.e. corresponding to 4.5 µg of starting RNA before fragmentation) were subjected to m6A-immunoprecipitation using $m^6A$ antibody-coupled protein A/G magnetic beads following the low/high salt procedure by Zeng and collaborators (Zeng et al. 2018). For each sample, 30 µl of protein A and 30 µl of protein G magnetic beads (Thermo Fisher Scientific, Waltham, MA, USA) were incubated on a magnetic rack and washed twice with IP buffer (150 mM NaCl, 10 mM Tris-HCl pH 7.5, IGEPAL 0.1 %, DEPC water) and resuspended in 500 µl of IP buffer. The magnetic beads were incubated overnight at 4 °C under gentle shaking with 5 µg of anti-$m^6A$ antibody (ABE572, Millipore, Burlington, MA, USA). Then magnetic beads were washed twice with IP buffer and resuspended in RNA mixture composed of 100 µl of 5X IP buffer, 44 µl of fragmented RNAs and 200 units of RNasin Plus Ribonuclease inhibitor (Promega, Madison, WI, USA) and the mixture was completed to 500 µl with IP buffer. The obtained MeRIP solution containing magnetics beads coupled to antibody and fragmented RNAs was incubated for 2 h at 4 °C with gentle shaking. After incubation, the MeRIP solution was incubated on a magnetic rack and the supernatant was removed. Beads were washed twice in IP buffer, twice in low salt buffer (50 mM NaCl, 10 mM Tris-HCl pH 7.5, 0.1 % IGEPAL, DEPC water) and twice in high salt buffer (500 mM NaCl, 10 mM Tris-HCl pH 7.5, 0.1 % IGEPAL, DEPC water) for 10 min at 4 °C under gentle shaking for each washing step. After extensive washing with IP buffer, RNAs hybridized to the antibody-coated magnetic beads were resuspended in 200 µl of RA1 buffer supplied in the Nucleospin RNA Clean up kit (Macherey-Nagel, France) and incubated 2 min at room temperature then 1 min on a magnetic rack. The supernatant containing immunoprecipitated RNA was collected and mixed with 400 µl of 100 % ethanol. Then the immunoprecpitated RNA was purified using affinity chromatography (Nucleospin RNA Clean up kit, Macherey Nagel, France) and eluted twice with 14 µl of RNase free water. These immunoprecipitated RNAs correspond to the immunoprecipitated fraction (IP).

**RT-qPCR:**

To validate the immunoprecipitation of $m^6A$ methylated RNA, a RT-qPCR was performed on immunoprecipitated RNA pools composed of equal amounts of RNA from each developmental stage for the two distinct experiments. Input and IP fractions were obtained as described above. The equivalent amount of 1,75 ng of Input and IP RNAs were used as starting template for the RT-qPCR protocol previously described (Riviere et al. 2011). Targets included one reference transcript (*ef1α*) and five conserved genes exhibiting distinct transcript $m^6A$-methylation in vertebrates (*c-myc*, *klf*, *mettl3*, *hnrnpa2b1* and *oct4*). Briefly, RNAs were reverse-transcribed using 200 U of M-MLV RT (Promega, Madison, WI, USA) and 100 ng random hexamers. Resulting cDNAs were assayed for target gene expression using the Input expression as reference. SYBR-green quantitative PCR was performed on a CFX96 apparatus (Bio-Rad, France). Gotaq qPCR master mix (Promega) was used in 40 cycles (95 °C/15 s, 60 °C/15 s) reactions. Target genes included one reference gene and 5 target genes with distinct transcript m6A methylation in vertebrates, and were amplified with the following primers: *Cg-ef1α* (CGI_10012474; forward: 5'-ACCACCCTGGTGAGATCAAG-3', reverse: 5'-ACGACGATCGCATTTCTCTT-3') as reference transcript of gene expression in *C.gigas* (Dheilly et al. 2011), genes described as methylated in vertebrates: *Cg-c-myc* (CGI_10002799; forward: 5'-CGGTCTCTCCCAAATTCTCCC -3', reverse: 5'-TGCTACTTCCACTTGCCCTG -3') (Huang et al. 2018; Batista et al. 2014) , *Cg-klf* (CGI_10019173; forward: 5'-GAAATCTCCGATGTTGCTGG-3', reverse: 5'-CTTTCCACGGTATTTGCGAG-3') (Batista et al. 2014), *Cg-mettl3* (gi762082209; forward: 5'-TGGAACCAAAGAAGAGTGTCAGA-3', reverse: 5'-AGAAATGAACAGTCTCCAAGGGA -3') (Coots et al. 2017) and *Cg-hnRNPA2B1* (gi762104361 ; forward: 5'-CCAGGGAGGCTACAATGAAGG -3', reverse: 5'-

ACACCACCACCAAAGCTGTT -3’) (Kwon et al. 2019) and a gene described as not methylated in vertebrates: *Cg-oct4* (XM_034479449; forward: 5’-GTGAAAGGTGCGCTAGAAAA-3’, reverse: 5’-GGACCACTTCTTTCTCCAGT-3’) (Batista et al. 2014). The methylation level was calculated as the normalized ratio of the IP and the Input signals by the formulae: 2^(-∆(Ct IP-Ct Input)).

The $m^6A$ motif was searched in the 1000 $m^6A$ peaks presenting the lowest FDR and the highest IP/Input fold change using Homer (v.4.10.4). The motif length was restricted to 5, 6 nucleotides. All peaks mapped on mRNAs and ncRNAs were used as the target sequences and the background sequences were constituted of 5% of the Input reads (pool of February development experiment sample) selected randomly using Samtools (v.1.9).

**Gene ontology analysis:**

The RNA sequences identified as differentially methylated across oyster development were identified using BlastN (Camacho et al. 2009; Cock et al. 2015; Altschul et al. 1997) against the GigaTON reference transcriptome database (Riviere et al. 2015) with default settings. Gene ontology (GO) analyses were carried out with the GO annotations obtained from the GigaTON database gene universe (Riviere et al. 2015). GO-term enrichment tests were performed using ClueGO plugin (v.2.5.7) (Bindea et al. 2009) on Cytoscape (v.3.8.0). The hypergeometric test with a FDR <5 % was used to consider significant GO term enrichment.

**Statistical analyses and graph production:**

The m$^6$A enrichment and the variation of the methylation level across oyster development was analyzed using one-way ANOVA (factor: development stage) followed by Bonferroni's post-hoc test when required, unless otherwise stated. Values were log-centered and reduced for heatmap production when stated. Methylation profiles were compared using Kolmogorov-Smirnoff tests (CDS regions were divided into 20 bins, start and stop codon intervals as well as 5' and 3' UTR regions were left unmodified from the default MetPeak output). Correlation between methylation level and transcript content across oyster development, as well as methylation variation and transcript content variation between stages were estimated using linear regression. A p-value <0.05 was considered significant. All bioinformatic analyses (unless stated otherwise) were performed using R (v.3.6.3) and RStudio (v.1.0.153) softwares. The R packages *eulerr* (Larsson 2019), *Complexheatmap* (Gu et al. 2016), *ggplot2* (Gómez-Rubio 2017), *Guitar* (Cui et al. 2016b), *PCATools* (Blighe and Lewis 2019), *corrplot* (Wei and Simko 2017) and Prism v.6 (Graphpad) software were used for figure production.
